# Brain Specific Estrogen Ameliorates Cognitive Effects of Surgical Menopause in Mice

**DOI:** 10.1101/2023.08.09.552687

**Authors:** Abigail E. Salinero, Charly Abi-Ghanem, Harini Venkataganesh, Avi Sura, Rachel M. Smith, Christina A. Thrasher, Richard D. Kelly, Katherine M. Hatcher, Vanessa NyBlom, Victoria Shamlian, Nyi-Rein Kyaw, Kasey M. Belanger, Olivia J. Gannon, Shannon B.Z. Stephens, Damian G. Zuloaga, Kristen L. Zuloaga

## Abstract

Menopause is a major endocrinological shift that leads to an increased vulnerability to the risk factors for cognitive impairment and dementia. This is thought to be due to the loss of circulating estrogens, which exert many potent neuroprotective effects in the brain. Systemic replacement of estrogen post-menopause has many limitations, including increased risk for estrogen-sensitive cancers. A more promising therapeutic approach therefore might be to deliver estrogen only to the brain thus limiting adverse peripheral side effects. We examined whether we could enhance cognitive performance by delivering estrogen exclusively to the brain in post-menopausal mice. We modeled surgical menopause via bilateral ovariectomy (OVX). We treated mice with the pro-drug 10β,17β-dihydroxyestra-1,4-dien-3-one (DHED), which can be administered systemically but is converted to 17β-estradiol only in the brain. Young (2.5-month) and middle-aged (11-month-old) female C57BL/6J mice received ovariectomy and a subcutaneous implant containing vehicle (cholesterol) or DHED. At 3.5 months old (young group) and 14.5 months old (middle-aged group), mice underwent behavior testing to assess memory. DHED did not significantly alter metabolic status in middle-aged, post-menopausal mice. In both young and middle-aged mice, the brain-specific estrogen DHED improved spatial memory. Additional testing in middle-aged mice also showed that DHED improved working and recognition memory. These promising results lay the foundation for future studies aimed at determining if this intervention is as efficacious in models of dementia that have comorbid risk factors.

## INTRODUCTION

The average woman spends ~1/3 of her life in menopause. By 2030, it is projected that ~14% of the entire world population will be menopausal women^1^. Menopause is an endocrinological transition, due to loss of ovarian hormones, that increases risk for cognitive decline and dementia^2-5^. In part this is through the exacerbation of dementia risk factors ^3-5^. For example, rates of type 2 diabetes increase among menopausal women^6-8^. Glucose metabolism within the brain decreases after the transition to menopause despite the brain’s high energy demands; this is accompanied by hippocampal volume loss, increased amyloid beta deposition, and impaired memory women^9-12^. Meta-analyses reveal that while early age at menopause is associated with an increased risk for developing dementia^13^, later age at menopause is associated with better cognitive performance and delayed cognitive decline^14,15^.

Loss of ovarian-produced estrogen during natural menopause or following surgical menopause (removal of both ovaries), is thought to contribute to the acceleration of cognitive decline. Estrogen exerts potent neuroprotective effects within the central nervous system (CNS)^16-18^. This includes maintaining energy balance through the regulation of brain glucose transport and metabolism^19,20^, enhancing blood brain barrier (BBB) integrity^21^, hastening the resolution of inflammation post-injury^22^, and regulating neuronal synaptic plasticity^23-26^. Estrogen receptors are expressed on neurons, glia, and other cell types such as endothelial cells throughout the brain^27-32^. Estrogen receptors are abundantly expressed within brain structures integral for learning and memory such as the hippocampus^33^ and cortex^31,34^. These structures are highly responsive to exogenous estrogen ^35,36^, which speaks to the major role estradiol plays in mediating brain homeostasis. Therefore, supplementing exogenous estrogens after menopause could help ameliorate post-menopausal cognitive decline.

Several large observational studies have shown that longer lifetime exposure to estrogen is also linked to better cognition and lower Alzheimer’s disease risk^37,38^. Unfortunately, clinical trials involving treatment with estrogen have been largely unsuccessful^39-41^. There is a new understanding that estradiol is likely only protective when it is given soon after menopause (<5 years)^42^ and in the right formulation (17β-estradiol rather than conjugated equine estrogens) for a long enough duration to preserve cognitive function^42,43^. Additionally, systemic treatment with estrogen has even been associated with adverse peripheral effects^39,44^ such as increased risk of estrogen-sensitive cancers^39^. Given these limitations, a more promising strategy may be to supplement estrogen exclusively to the brain. In 2015, Prokai *et. al* characterized a novel estrogen prodrug, 10β,17β-dihydroxyestra-1,4-dien-3-one (DHED). When administered systemically to mice or rats, DHED is converted to 17β-estradiol only in the brain while remaining inert throughout the body^45^. *In vitro*, DHED does not enhance estrogen-sensitive tumor size ^45^. Subsequent studies have demonstrated that DHED ameliorates some of the adverse effects of menopause such as hot flashes^46^ and depressive symptoms in rodents^47^. These significant studies demonstrate the potential of DHED as a brain-specific estrogen delivery option. However, DHED’s potential to improve cognition has been understudied, and it has not been examined in middle-aged mice. Middle age itself is a time in which chronological aging processes converge with the menopausal transition to contribute to the increased vulnerability to cognitive decline^48^. In this study, we sought to validate that long-term DHED delivery is efficacious during this critical aging timepoint of middle-age.

Our main goal was to evaluate whether DHED could improve cognition in post-menopausal middle-aged female mice. We used bilateral ovariectomy to model surgical menopause and delivered estrogen to the brain via silastic capsule implant of DHED in both young and middle-aged mice. We measured cognition after long-term treatment through a series of behavioral tests. We identified cognitive improvements in mice of both ages, further supporting the therapeutic potential for DHED.

## METHODS

### ANIMALS AND EXPERIMENTAL DESIGN

All experiments were approved by the Albany Medical College Animal Care and Use Committee and in compliance with the ARRIVE guidelines.

#### Study 1 Timeline

Female C57BL/6J mice were purchased at 9 weeks of age from Jackson Laboratories (Bar Harbor, ME, USA). Upon arrival, mice were randomly assigned to cages [group housed (3-5 per cage) at 70-72 °F, 30-70% humidity, with a 12 h light/12 h dark cycle (7 a.m. on/7 p.m. off)]. Mice were group housed for the duration of the study. Mice were provided with normal chow diet (Purina Lab Diet 5P76) and water ad libitum. Following one week of acclimation, cages of mice were randomized to treatment groups based on cage ID number and received bilateral ovariectomy and a intrascapular subcutaneous silastic capsule implant containing either DHED (mixed to a 1:10 ratio with cholesterol packed to a length of 4mm) or vehicle (cholesterol only packed to a length of 4mm). One month later, mice underwent Morris water maze (protocol described below) to assess spatial learning and memory. The following week mice were deeply anesthetized with isoflurane and intracardially perfused with saline and tissues were collected. Experiments were conducted in 1 cohort of 16 mice (N=8 mice per group). Experimenters were blinded to DHED status.

#### Study 2 Timeline

Female C57BL/6J mice were purchased at 44 weeks of age from Jackson Laboratories (Bar Harbor, ME, USA). Acclimation, housing conditions, and randomization to treatment group were the same as described for study 1. After 1 week of acclimation, mice received bilateral ovariectomy and a silastic capsule implant containing DHED in a ratio of 1:30 (low dose), 1:10 (medium dose), 1:3 (high dose), or vehicle (cholesterol), each packed to a length of 4mm. A reference group of 10 age-matched females received a sham surgery and vehicle implant. Capsule implants were replaced every 6 weeks. At 3.5 months after ovariectomy, mice underwent a series of behavior tests to assess cognitive deficits. Next, a glucose tolerance test was performed to assess metabolic status. Three and a half months after surgery, mice underwent behavior testing (detailed below) in the following order: Anxiety-like behavior was assessed using the marble burying test, anxiety-like behavior and general locomotor activity were tested with the open field. Spatial recognition memory and episodic-like memory were assessed in the object place recognition test (OPRT) and novel object recognition test (NORT). Working memory was assessed in the Y maze, and activities of daily living were assessed with the nest building test. For all behavior tests mice were allowed to acclimate to the behavior room for one hour prior to testing in the same ambient conditions used during the test. Lights were kept dim except during marble burying, which took place under fluorescent lights (lux=~400). Open field, OPRT, NORT, and Y maze were video recorded and analyzed using automated tracking software (ANY-maze 5.1, Stoelting, Wood Dale, IL).The following week mice were deeply anesthetized with isoflurane and intracardially perfused with saline and tissues were collected. Mice were group housed for the duration of the study. Experiments were conducted in 3 cohorts of 16-17 mice (n=9-10 mice per group). Final n numbers for each figure are noted in figure legends. Experimenters were blinded to DHED status.

#### Bilateral Ovariectomy (OVX)

Mice were anesthetized with isoflurane and body temperature was maintained. An incision was made to the caudal abdomen and underlying peritoneum. The first ovary was then located and pulled out before ligation distal to the uterus. A cut was made above the oviduct to remove the ovary. Once hemostasis was verified, the uterus was returned into the abdomen. The second ovary was removed in the same manner. After the removal of both ovaries, incisions were closed using 4-0 silk sutures and tissue adhesive. For age-matched intact mice (study 2), the abdomen and peritoneum were incised, and the ovaries were identified and pulled out. The ovaries were then placed back into the abdomen without ligation or excision. Wound clips were placed on the incisions. Mice were allowed to recover from anesthesia before returning to their home cage. Buprenorphine (100 µl of 0.03mg/mL) was administered via subcutaneous injection (2x day for 3 days) for pain management.

#### Preparation of Silastic Capsules

Silastic capsules were prepared 3 days prior to implantation. On day 1, 10mm of silastic tubing (ID: 1.47mm OD: 1.96mm; 508-006 Dow Corning) was cut and one side was clogged with ~2mm of silicone glue (Permatex #80050). This was allowed to set overnight protected from dust. The next day, DHED powder (SML1642-25MG, Millipore) was mixed in a 1:10 ratio (study 1) with cholesterol (45-C8667, Krackeler Scientific) and silastic tubing was packed to 4mm with the DHED mixture or cholesterol only (vehicle). For study 2, to assess the efficacy of different ratios of DHED, capsules were packed with DHED powder in ratios of 1:30, 1:10, 1:3, or cholesterol only (vehicle). The remainder of the tube was sealed with silicone glue and allowed to set overnight. On day 3, the capsules were washed in 70% ethanol and soaked overnight in sterile saline.

#### Silastic Capsule Implant

Immediately following ovariectomy, while mice were still anesthetized with isoflurane, a small incision was made between the shoulder blades of the mice and the capsule was inserted subcutaneously with sterile forceps. Incisions were closed using 4-0 silk sutures and tissue adhesive. For study 1, capsules remained in place for the duration of the study (4 weeks). For study 2, capsules were initially implanted immediately after OVX as described for study 1 and replaced every 6 weeks to ensure continuous administration of the drug. During implant replacement, the old capsule was removed using sterile forceps before the new one was placed. Buprenorphine (100 µl of 0.03mg/mL) was administered via subcutaneous injection (2x day for 3 days) for pain management.

### BEHAVIOR

#### Morris Water Maze

Morris Water Maze (MWM) was performed as previously described^49^. Mice were placed in a pool 125 cm in diameter. Mice were trained to swim to a platform submerged 2.5cm under water made opaque with white tempera paint. The temperature of the pool was 20 to 22 degrees Celsius. MWM took place in three different types of trials: visible, hidden, and probe. During the visible trials (day 1), mice were trained to locate the escape platform submerged below the surface of the water with the aid of visual cues. Mice underwent 6 visible trials, separated by a 30-minute inter-trial interval each. Mice were carefully placed near the wall of the pool in two alternating locations. The trial ended when the mouse located the platform. Mice were rescued and placed in a recovery cage under a heat lamp. If the mouse failed to locate the platform after 3 minutes, they were guided to the platform by placing a finger on the platform. If they still did not locate the platform, they were brought to the platform by being gently dragged by the tail. This was repeated until they stayed on the platform for at least 10 seconds. Throughout the visible trials, the platform was not moved. The following day, mice were trained to locate a hidden platform, using extra-maze cues for spatial reference and orientation (hidden trials; day 2). Visual cues were placed on the walls of the pool above the water line at the borders of the quadrants. The platform was returned to the same place and remained in the same location for all hidden trials. There were 6 hidden trials with a 30-minute inter-trial-interval. Twenty-four hours after the final hidden trial, spatial memory retention was assessed through a three-minute probe trial (in which the platform was removed; day 3). For the visible and hidden trials, latency to find the platform over the course of the trials was measured. For the probe trial, the straightness of the path used to reach the target location was measured (path efficiency). All trials were video recorded and analyzed using automated tracking software (ANY-maze 5.1, Stoelting, Wood Dale, IL).

#### Marble Burying

Allentown cages were filled with wood chip bedding to a depth of 1 inch. 12 identical black marbles were gently placed on top of the bedding in 4 rows of 3 marbles. One mouse was placed in the corner of each cage. The cages were covered with a filter top and the mice were undisturbed for 30 minutes. Mice did not have access to food or water during the test. After 30 minutes, the mice were removed carefully to not move the marbles and placed back in their home cages. Pictures were taken from an aerial view of the cages. Marbles were cleaned with 70% ethanol to prevent odor cues between mice. Three independent scorers blinded to treatment counted the number of marbles buried in each cage based on the picture. Marbles were considered buried if two-thirds or more of their surface area was covered by the bedding. The average of the three scores was taken as the number of marbles buried.

#### Open field

As previously described^49,50^, mice were placed in a 49.5 × 49.5 cm square arena and allowed to explore freely for 10 min, then removed and placed in a “recovery cage” so as not to expose them to naïve cage mates. Anxiety-like behavior was measured as the amount of time spent in the center of the arena. General locomotor behavior was assessed as the measure of total distance traveled (pathlength).

#### Object Place and Novel Object Recognition

The combined test took place as three 10-minute trials: a “training” trial, the “OPRT” trial and the “NORT” trial, with an inter-trial interval of 1hr. The open field arena was used, with a horizontal piece of black tape on the north wall serving as a visual cue. In the training trial, mice were placed in the open field arena and allowed to explore two identical objects, one in the northwest (NW) corner and one in the northeast (NE) corner, each 8 cm from the wall. After the training trial, mice were placed in a recovery cage for 2 hours, and the open field box and objects were cleaned with 70% ethanol. In the OPRT test trial, the NW object was placed in its original position while the NE object was moved to the southwest (SW) corner. Mice were allowed to explore again for 10 minutes before being returned to their recovery cage for 2 hours. In the NORT test trial, the NW object was returned to its original position in the arena, along with a novel object in the SW corner. Mice were allowed to explore again for 10 minutes. For the training trial, preference for the NE object vs NW object was computed as [(NE object time)/(NE + NW object time) x 100]. For the OPRT and NORT, recognition indexes [(SW object time)/(SW + NW object time) x 100] were computed. To evaluate spatial recognition memory and episodic-like memory, the difference between the recognition indexes and the preference during the training trial were calculated. Exclusion criteria included failure to explore both objects as well as exploring both objects for less than 10 seconds during the training trial. Additionally, animals were excluded if they explored objects for less than 4 seconds during the NORT or OPRT. 9 animals total were excluded from OPRT and 10 animals were excluded from NORT based on these criteria.

#### Y Maze Spontaneous Alternations

Mice were placed in a Y-maze (Stoelting, Wood Dale, IL) with 3 identical arms (5 cm W x 35 cm L x 10 cm H) and allowed to explore freely for 3 minutes. Data from the first minute was analyzed. Arm entries were recorded. Each set of 3 non-repeating arm entries was counted as a triad. Working memory was measured as the % of spontaneous alternations [(# of triads)/(# arm entries – 2) x 100]. Mice with fewer than 4 arm entries were excluded from the test (1 mouse was excluded based on this criteria).

#### Nest Building

Nest building took place as previously described^49,51^. Mice were individually housed overnight and were provided intact nestlet material. 16 hours later, mice were carefully removed from the cages and group-housed in their home cage. The nests were scored by three independent scorers, blinded to conditions. Nests were scored on a scale of 1-5 in 0.5 point increments ^52^, with 1 being the lowest possible score and 5 being the highest. Scores were averaged between the three scorers.

#### Glucose Tolerance Test

Glucose tolerance test was carried out as previously described^49,50,53^. After the completion of behavior testing, mice were fasted for 16 hours overnight, and the next morning baseline blood glucose levels in saphenous vein blood were measured by glucometer (Verio IQ, OneTouch, Sunnyvale CA, USA). Each mouse received 2g/kg of glucose via i.p. injection and blood glucose levels were re-measured at 15, 30, 60, 90, and 120 minutes post-injection.

#### Quantitative PCR

qPCR was performed according to our previously published methods ^49,50^. Flash frozen hippocampi were thawed and homogenized in 50µL RNA-Later (45-R0901-100MLsigma). RNA was extracted from 25uL of homogenate using the RNeasy® Plus Mini Kit (Qiagen, Catalog number 74134). RNA concentrations were determined using ThermoScientific NanoDrop One and RNA was converted to cDNA using a High-Capacity cDNA Reverse Transcription Kit (Applied Biosystems, Catalog number: 4368814). The qPCR reactions were performed using TaqMan Gene Expression Master Mix (Applied Biosystems, Catalog number 4369016) in the presence of TaqMan Assays with primer/probes for ERα (Mm00433149_m1), ERβ (Mm00599821_m1) and GPER1 (Mm02620446_s1) as target genes. RPL13A (Mm05910660_g1) was used as the housekeeping gene. Data is represented as fold change relative normalized expression compared to OVX-vehicle-treated mice using Bio-Rad CFX Maestro software.

#### Statistics

Data were analyzed with Prism 8.1 (GraphPad Software, San Diego, CA, USA). A student’s T-test was used when comparing 2 groups only (e.g., Morris water maze probe trial in Study 1). For study 2, one-way ANOVA with Dunnett’s correction for multiple comparisons was used to compare all groups to OVX-vehicle. Gonadally intact vehicle treated mice are only represented for reference. For behavior data in which performance is judged relative to chance (e.g recognition index), data are analyzed with a one-sample T-test vs. chance. For data that was tracked over time (e.g. Morris water maze escape latency and weight change), mixed effects ANOVAs [DHED X trial (1-12)] were used. The robust regression and outlier removal test (Prism 8.1) was used. Data is presented as mean ±SEM on bar graphs and mean +/-SEM on line graphs.

## RESULTS

### DHED improves spatial learning and memory in ovariectomized young mice

To investigate whether DHED could improve cognition when administered long term via silastic capsule, young (2.5 months old) mice were given a bilateral ovariectomy (OVX) and implanted with a subcutaneous silastic capsule of either vehicle (cholesterol) or DHED (ratio of 1:10 in cholesterol). One month later, spatial learning and memory were assessed through the Morris water maze (**Figure 1A**). Across repeated visible trials, when the platform was marked by a flag, both groups significantly reduced their time to reach the platform, and no group differences were seen in this latency (mixed effects ANOVA, main effect of trial, p<0.0001, trial x treatment, p=0.4857), indicating that both groups demonstrated cued learning equally well. However, when the flag was removed and animals had to rely on spatial cues in and around the maze to navigate to the submerged platform (hidden trials), escape latency was significantly better in DHED treated animals (mixed effects ANOVA, main effect of trial, p=0.0363, trial x treatment, p=0.0.0116; **Figure 1B**). When spatial memory was assessed in the probe trial, DHED treated mice showed a significantly better path efficiency to locate the area where the platform had been (**Figure 1C, 1D**; Student’s T-test p=0.0265). These data indicate that DHED significantly improves spatial learning and memory in a surgical menopause model.

**Figure 1.**
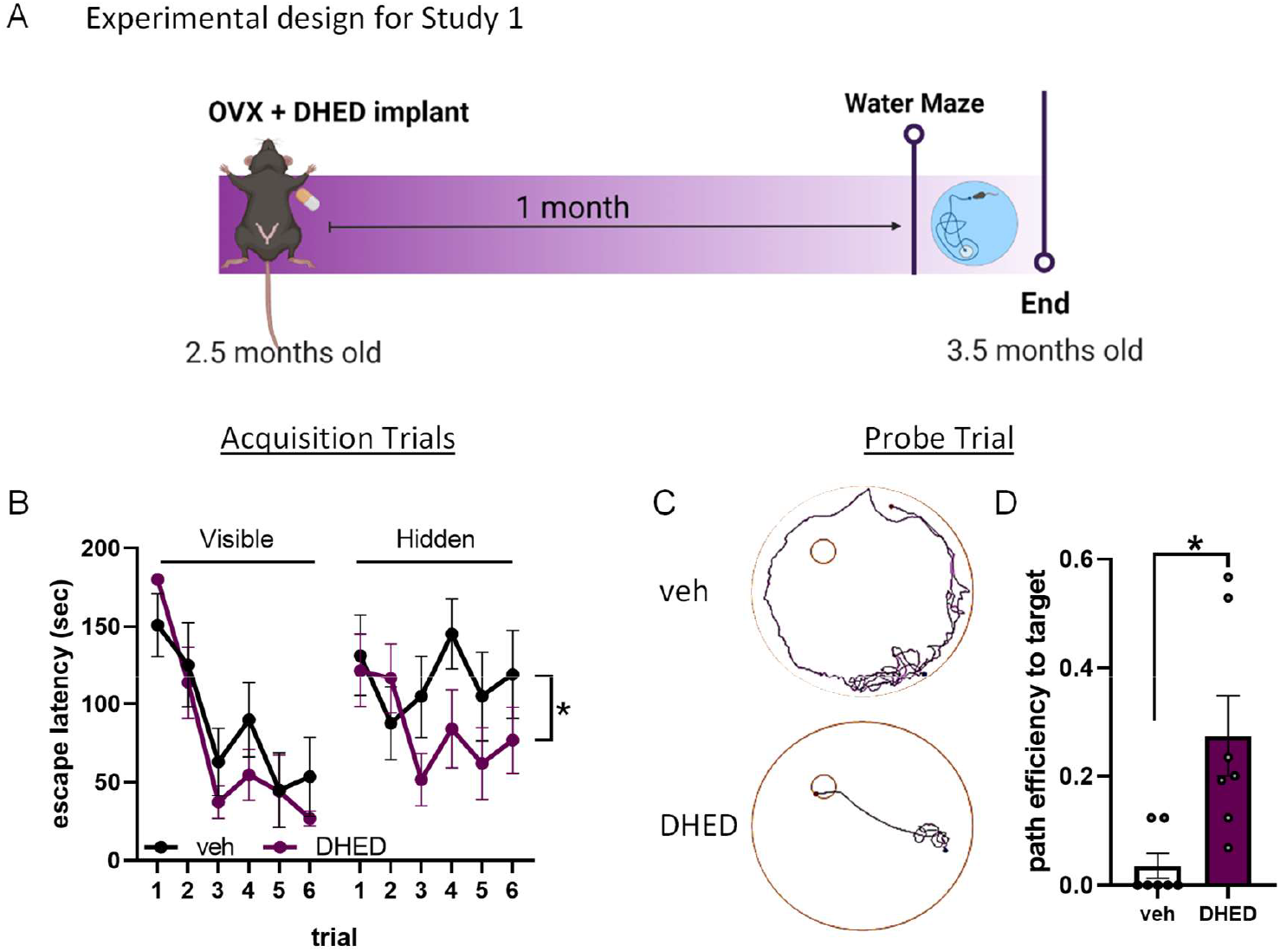
DHED improves spatial learning and memory in ovariectomized young mice. (A) Experimental timeline for Study 1. Figure created with BioRender. (B) Escape latency during the visible (left) and hidden (right) trials of Morris water maze (MWM) performed one month after ovx and vehicle (veh) / DHED implant. Mixed effects ANOVA showed a trial x treatment interaction for hidden trials only *p=0.0116. (C) Representative plots of the path vehicle-treated mice (top) and DHED-treated mice (bottom) used to locate the area where the escape platform had been during the probe trial. (D) Path efficiency to reach the area that the escape platform had been during the probe trial. N=7/group. Student’s *t* test *p=0.0172. Data are presented as mean ±SEM.

### DHED does not exacerbate metabolic effects of ovariectomy in middle-aged females

Loss of estrogens during menopausal transition has been shown to increase metabolic burden in women, such as increased visceral adiposity and glucose intolerance^54-57^. To investigate metabolic effects in a more clinically relevant age group, we performed bilateral ovariectomy in middle-aged (11 months old; **Figure 2**) females and treated them with silastic capsules containing either vehicle (cholesterol), DHED in a ratio of 1:30 (low dose), 1:10 (medium dose), or 1:3 (high dose). Capsules were replaced every six weeks to ensure continuous drug supply for the 3.5 month duration. An ovary-intact vehicle treated group was included for reference purposes on each graph (green line = mean, shading = standard error). Weight was tracked for the duration of the study (**Figure 3A**). Percent change in body weight at the end of the study relative to the mouse’s starting weight was calculated (**Figure 3B**). There were no group differences in weight gain (mixed effects ANOVA time x treatment p=0.7519; one way ANOVA p=0.7278). Visceral adipose tissue, which is known to increase in humans after menopause onset and is associated with increased risk for cardiovascular disease and other metabolic diseases, was collected at the end of the study and weighed (**Figure 3C**). There were no group differences in visceral adipose tissue as a % of body weight (one way ANOVA p=0.8889). Glucose intolerance was assessed through a glucose tolerance test. Overall, there was a time x treatment interaction (mixed effects ANOVA p=0.0412). Specifically, compared to vehicle-treated mice, mice that received the high dose DHED showed significantly higher blood glucose levels at 60 and 90 minutes after glucose injection (**Figure 3D**; Dunnett’s multiple comparison test: 60 min p<0.0008, 90 min p=0.0236 DHED high vs vehicle). While no significant group differences were observed when analyzing the area under the curve (AUC) for the glucose tolerance test (**Figure 3E**; one way ANOVA p=0.1018), there was a trend for the high dose DHED-treated animals to have greater AUC than vehicle treated animals (p=0.0535). These data indicate that at low-to-moderate doses DHED does not impair glucose tolerance. Uterine weight fluctuates with variations in levels of circulating estrogen and can be used as a proxy measurement of circulating estrogens. Uteri were dissected at the end of the study and wet weights taken (**Figure 3F**). As expected, ovariectomized mice had lower uterine weights than the ovary-intact reference group and there were no group differences in uterine weight among the vehicle and DHED-treated mice (one way ANOVA p=0.1363). Importantly, this data indicates that DHED does not exert estrogenic effects on this estrogen sensitive peripheral organ. Overall, these data suggest that low and medium doses of DHED do not affect metabolic status of menopausal mice.

**Figure 2.**
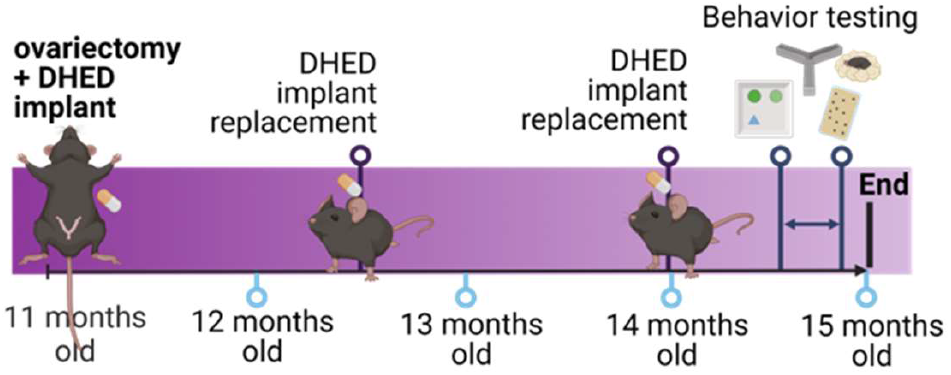
Experimental design for study 2. Mice were ovariectomized and implanted with DHED or vehicle at 11 months of age. Capsules were replaced every 6 weeks to ensure continuous supply of the drug. Behavioral assays were performed 3.5 months after ovariectomy/implant. Figure created with BioRender.

**Figure 3.**
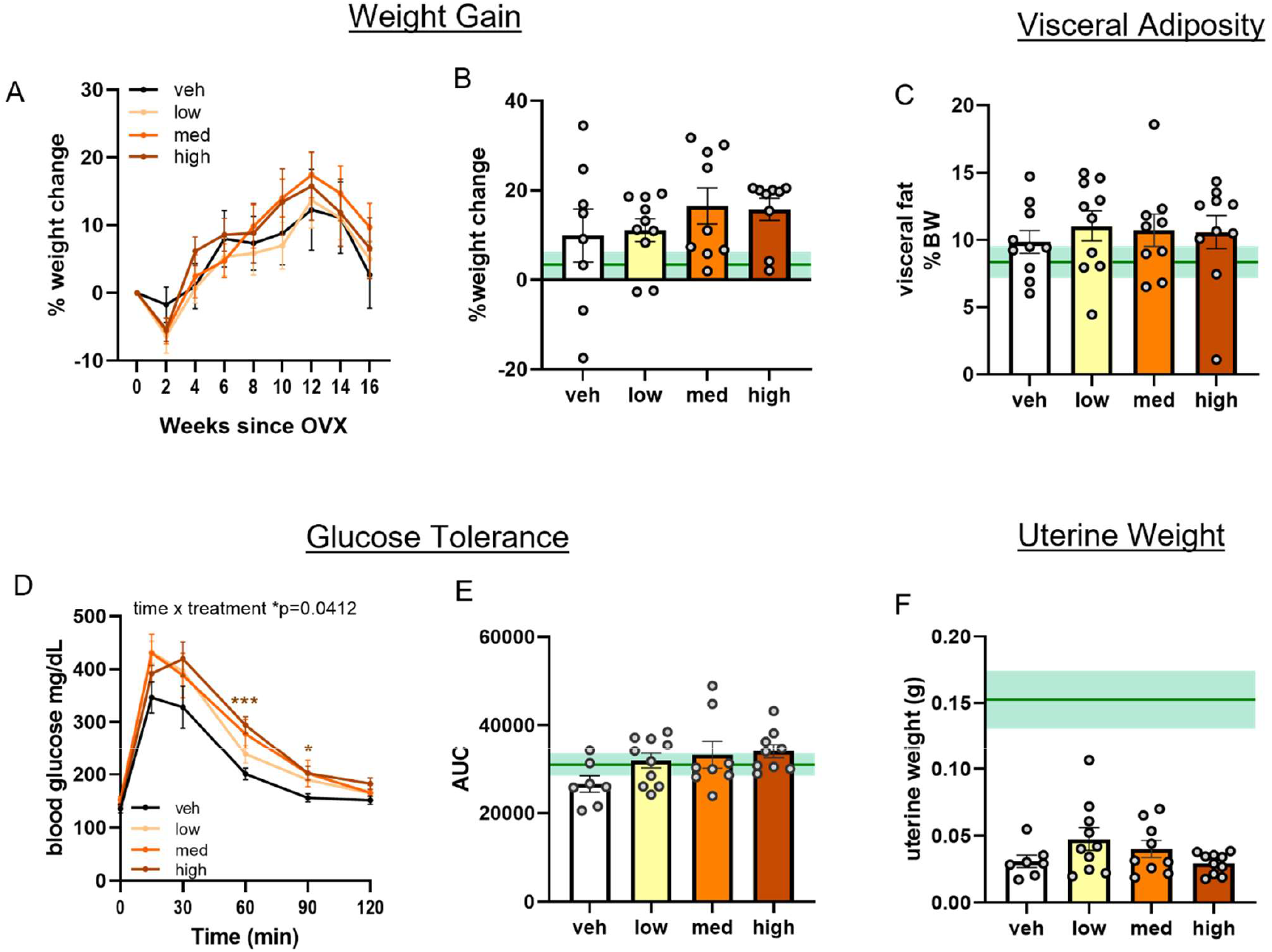
DHED does not exacerbate post-menopausal metabolic effects in middle-aged females. (A) % weight change over the course of the study. (B) Weight change (%) from start to end of study. (C) Fat pad weight as % body weight at the end of the study. (D) Blood glucose concentrations (mg/dL) over 2 hours post-glucose challenge. Mixed effects ANOVA time x treatment interaction *p=0.0412 with Dunnett’s post hoc test vs vehicle *p<0.05, ***p<0.001. (E) Area under the curve (AUC) from the glucose tolerance test. (F) Uterus weight at the end of the study. N=7-10/group. One-way ANOVA with Dunnett’s post hoc test vs vehicle. Data are presented as mean ±SEM. Gonadally intact, vehicle treated, age-matched reference group represented on bar graphs as mean (dark green line) ±SEM (light green shading). veh=vehicle; med=medium.

### DHED improves working memory and spatial recognition memory in ovariectomized middle-aged females

To evaluate the ability of DHED to improve memory in middle-aged female mice using a surgical menopause model (OVX), we performed a series of behavior tests 3 months (age 15 months) after OVX and vehicle or DHED treatment. To assess working memory, mice were placed in a Y-maze and allowed to freely explore (**Figure 4A**). Spontaneous alternations (%) were analyzed as a measure of working memory (**Figure 4B**). Compared to vehicle-treated mice, mice that received either a medium or high dose of DHED showed significantly higher % alternations, indicating a better working memory (one way ANOVA, p=0.0147; vehicle vs DHED medium p=0.0362; vehicle vs DHED high p=0.0215). Mice that received a low dose of DHED did not perform significantly better than vehicle treated mice (p=0.8334). There were no group differences in overall number of arm entries (**Figure 4C**), indicating that differences in alternations were not related to overall exploratory differences (one way ANOVA p=3168). Taken together, these data indicate that while long term treatment with low dose DHED is not sufficient to improve working memory, treatment with DHED at the medium and high doses improved working memory in ovariectomized middle-aged females. We assessed spatial recognition memory through an object place recognition test (OPR; **Figure 4D**). OPR was calculated as the difference in preference for an object during the training trial (old place) vs the preference for same object during the test trial (new place). Compared to vehicle-treated mice, mice that received either a medium or high dose of DHED showed significantly greater preference during the test trial (**Figure 4E**), indicating a better object place memory (one way ANOVA p=0.0363; vehicle vs DHED medium p=0.0465; vehicle vs DHED high p=0.0252). Mice that received a low dose of DHED did not perform significantly better than vehicle (p=0.2848). No group differences were observed in total object exploration time (**Figure 4F**; one way ANOVA p=0.7135) suggesting that the differences observed during the OPR task were not related to differences in exploration.

**Figure 4.**
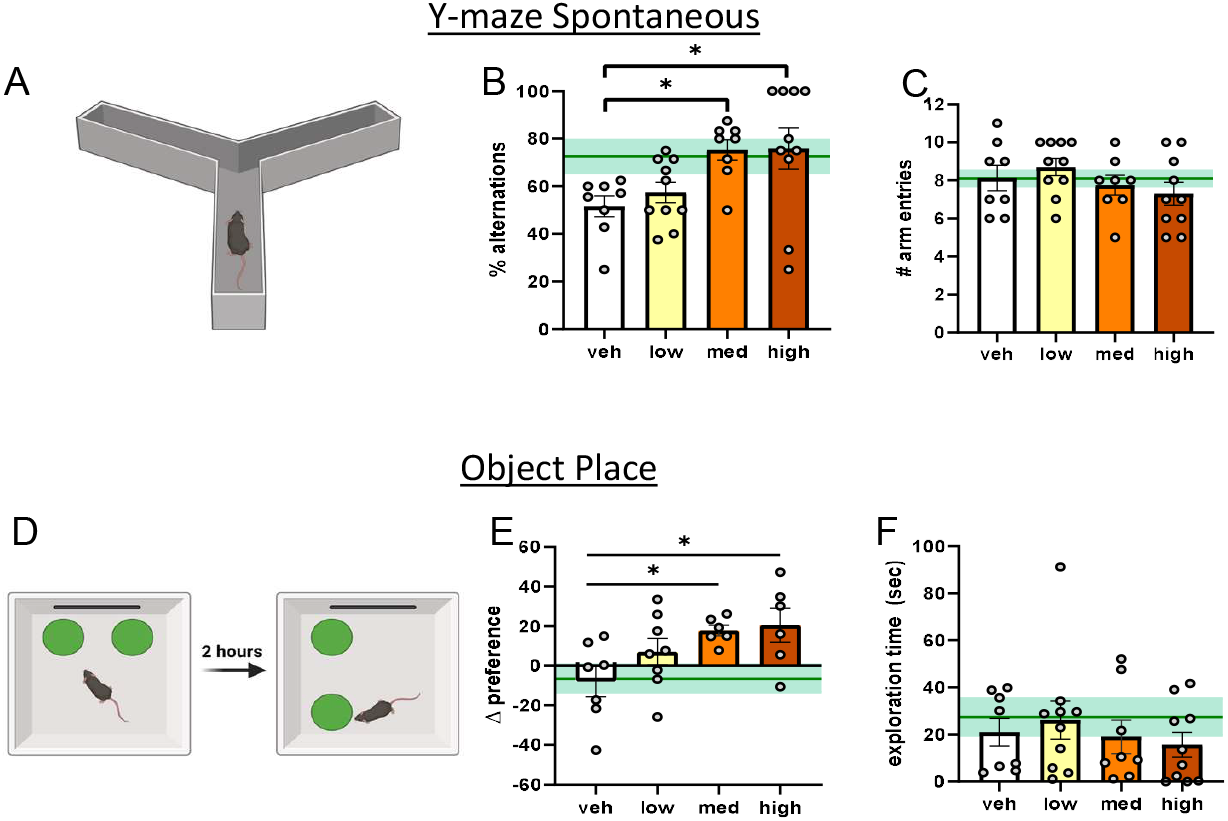
DHED improves working memory and spatial recognition memory in post-menopausal middle-aged females. (A) Schematic of Y maze spontaneous alternations (created in BioRender). (B) % alternations (C) Number of arm entries. N=8-10/group. (D) Schematic of object place recognition (created in BioRender) depicting the training and testing trial (E) Difference between % time spent with object in its new place during testing trial vs % time with that same object during training trial (F) Total exploration time with both objects. N=6-8/group. Data are presented as mean ±SEM. One-way ANOVA with Dunnett’s post hoc test vs vehicle *p<0.05. Gonadally intact, vehicle treated, age-matched reference group represented on bar graphs as mean (dark green line) ±SEM (light green shading). veh=vehicle; med=medium.

We assessed episodic-like memory in a novel object recognition test (NOR) after a two-hour inter-trial interval immediately following object place recognition test (**Sup. Figure 1A-B**). We did not detect any group differences via one way ANOVA (p=0.0899). However, secondary analysis (one-sample t-test vs 0, chance) revealed that the medium DHED group showed a significant preference for the novel object (p=0.0461) while the other groups did not. This indicates that while OVX mice are impaired in this test of episodic-like memory, mice treated with a medium dose of DHED after OVX have preserved episodic like-memory.

Since cognitive tests can be influenced by activity levels, we evaluated locomotor activity in the open field test and found no differences (**Sup. Figure 1C**; one way ANOVA p=2861). Anxiety-like behavior can also influence cognitive performance. We assessed anxiety-like behavior via the % of the time spent in the center of the open field (**Sup. Figure 1D**) and via defensive marble burying behavior (**Sup. Figure 1E**). We did not observe any significant differences in either test. Activities of daily living were assessed through a nest building test (**Sup. Figure 1F**), however, no group differences were observed. Overall, these data suggest that anxiety-like behavior and activities of daily living are not worsened by DHED. This further supports that the DHED-related improvements in memory tasks we report above are not due to confounding influences such as anxiety. Overall, these data suggest that long term treatment of DHED has beneficial effects on several cognitive aspects including working and spatial recognition memory.

### DHED does not change hippocampal expression of estrogen receptor mRNA

Given the improvements in hippocampal-dependent learning and memory tasks we observed in DHED treated mice, we sought to examine whether long term treatment with DHED altered expression of estrogen receptor genes in the hippocampus of our middle-aged mice (**Figure 5**). We did not observe any changes in hippocampal mRNA expression of estrogen receptors (ER) α or β (one way ANOVA p=0.7709 for ERα; p=0.5829 for ERβ; **Figure 5A and 5B**). The ratio of ERα to ERβ has been reported to decrease in the brain with age^58^ and to be functionally associated with increased synaptogenesis and transcription of synaptic plasticity genes ^59^. When we calculated the ratio of ERα to ERβ mRNA we did not observe any group differences (**Figure 5C**; one way ANOVA p=0.2025) We also did not observe any differences in G-protein coupled estrogen receptor 1 (GPER) (p=0.6061; **Figure 5D**). These data show that DHED treatment does not alter hippocampal expression of estrogen receptors genes.

**Figure 5.**
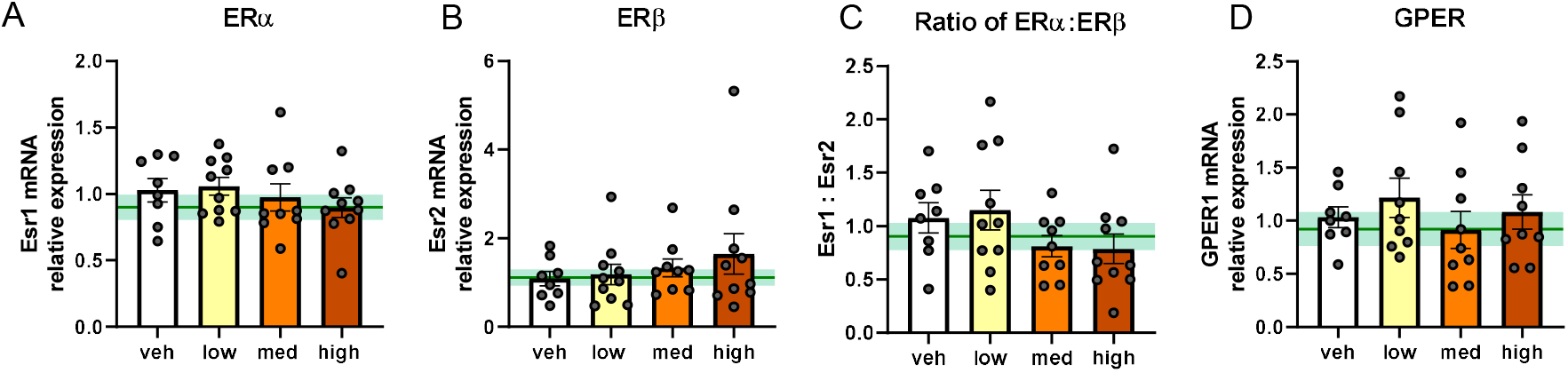
DHED does not change hippocampal expression of estrogen receptor mRNA. Relative expression of mRNA of estrogen receptors: ERα (Esr1) (A), ERβ (Esr2) (B) The ratio of ERα:ERβ (C), and GPER (GPER1) (D). mRNA levels were quantified using the ΔΔCq method and reported as fold change relative to vehicle-treated (veh) females. N=8-10/group. Data are represented as mean ±SEM. One-way ANOVA with Dunnett’s post hoc test vs. vehicle. Gonadally intact, vehicle-treated age-matched reference group represented on bar graphs as mean (dark green line) ±SEM (light green shading). med=medium.

## DISCUSSION

The goal of this study was to determine if the estradiol prodrug DHED can be administered long term via a silastic capsule implant to rescue cognitive deficits after surgical menopause. Others have previously demonstrated that DHED rapidly converts to estradiol in the brain^45^. However, whether cognitive performance can be improved by silastic capsules of DHED over 3.5 months had yet to be assessed. In our current study, we aimed to improve cognition in middle-aged females with surgical menopause by treating them with a subcutaneous silastic capsule implant of DHED. Overall, we found that DHED did not significantly alter metabolic status in middle-aged, post-menopausal mice. In both young and middle-aged mice, DHED improved spatial memory, and in middle-aged mice DHED also improved working memory. These results lay the groundwork for a novel therapeutic intervention for cognitive decline in women without increasing risk for adverse peripheral effects of traditional estrogen replacement therapy.

Systemic treatment with estrogen has been associated with adverse peripheral effect ^39,44^ such as increased risk of estrogen sensitive cancers^39^. This could be due to the direct pro-oncologic effect of estrogen on these peripheral tissues. By restricting estrogen’s actions to the brain, DHED represents a more promising opportunity for therapeutic intervention. In this study we found that DHED did not alter uterine wet weight, weight gain, visceral adiposity, or glucose tolerance in middle-aged females. Uterus wet weight is a proxy measurement for circulating estrogens. Our finding that DHED does not alter uterus weight indicates that DHED does not exert estrogenic effects on the uterus. This is consistent with what others have reported regarding uterus weight in response to DHED ^45,60^. Ours is the first study to examine the effect of DHED on metabolic measures. Systemic 17β-estradiol treatment reduces weight gain after OVX in middle-aged rats^61 61^ as well as young OVX mice^62^. This is associated with a reduction in food intake^62^. Perhaps our finding that DHED does not alter weight is due to the lack of peripheral 17β-estradiol in these mice. In the periphery, estrogen activates metabolic mediators such as insulin, leptin, and GLP-1^63^. Although some studies have linked the anorexigenic effects of systemic 17β-estradiol to its signaling within the hypothalamus, direct administration of 17β-estradiol to the hypothalamus has shown mixed results ^64,65^. We are the first to demonstrate that DHED does not have adverse metabolic consequences as assessed by body weight, glucose intolerance, and visceral adiposity.

In this study we found that in young ovariectomized mice, DHED improves spatial learning and memory. Specifically, we found that while mice are equally able to learn the association between swimming to the flagged target and being rescued from the water maze (associative learning), DHED improves cued learning as measured by the trials in which the platform is hidden, and mice must use spatial cues around the maze and room to locate the platform. In naturally cycling females, Morris water maze performance and search strategy vary depending on estrogen levels throughout the estrous cycle, with phases of high estrogen being associated with better performance^66,67^. One group assessed the effect of DHED on spatial memory using a different behavior test (radial arm maze) in an APPswe/PS1dE9 transgenic Alzheimer’s disease (AD) mouse model^60^. Among female AD mice, regardless of ovariectomy, DHED improved spatial memory. However, our study is the first to assess the effect of DHED on spatial recognition memory and to take into consideration the important variables of age, DHED dosage, and longerterm treatment which are likely to be clinically important variables given that age and dosage alters the efficacy and long term outcomes of HRT in women. In middle-aged females, we found that DHED significantly improved spatial recognition memory in the OPR test. Though no other study has examined DHED’s effect on spatial recognition memory, there is robust data on the ability of exogenous estrogen to enhance object place performance in rodents^68-70^. Inhibition of local E2 synthesis within the hippocampus impairs object recognition in ovariectomized female mice^71^. Overall, our data indicates that DHED has a similar beneficial effect of spatial recognition as estrogen in ovariectomized mice.

We found that in middle-aged females, DHED in a ratio of 1:10 or 1:3 improves working memory. To assess working memory in this study we used Y maze spontaneous alternations paradigm. Currently, only one other study has assessed the effect of DHED on working memory^45^. Prokai *et al* performed a delayed match-to-sample plus maze test in rats after ~1.5 months of DHED treatment (via alzet mini-pump)^45^. The authors found that DHED significantly improved working memory (reduction in the number of errors). In this study we found that mice treated with DHED over a longer time frame (3.5 months) exhibit enhanced working memory via Y maze spontaneous alternations paradigm. Though further investigation would be helpful, our work along with that of Prokai *et al* provides evidence of DHED’s ability to improve working memory across multiple behavioral assays and in multiple species over the long term.

Given that DHED is an estradiol pro-drug, we sought to establish whether estrogen receptor expression is altered by DHED administration. Previous studies have shown that inhibition of estrogen receptors (ER’s) ERα or ERβ in the hippocampus impairs object place recognition and novel object recognition in ovariectomized mice^72^. When we assessed levels of estrogen receptor mRNA in the hippocampus of middle-aged, DHED-treated mice, we found that ERα, ERβ as well as GPER were all similar to levels of intact, age-matched mice. Future studies are needed to determine which estrogen receptor(s) are mediating the protective effects of DHED. Improving our understanding of the mechanism by which DHED improves cognition is critical. It is highly likely that the mechanism is the same as that of estrogen in the brain. In this regard, so far data suggests that E2-mediated memory improvement occurs through estrogen receptors and depends on numerous cell signaling cascades such as ERK, PI3K/Akt, and mTOR as well as epigenetic processes such as DNA methylation^17,24,68,71-73^. Though outside the scope of this investigation, future research is needed to confirm that DHED administration results in upregulation of these same cascades.

Systemic hormone replacement therapy has shown mixed results in ameliorating menopausal cognitive decline^39-41^. Growing data suggests that these varied results may be due to the length of time between menopause onset and the initiation of estrogen replacement therapy, the “timing hypothesis”. Rodent data supports that there is a critical window after ovariectomy in which estrogen replacement therapy is beneficial, which is not related to chronological age^74,75^. However, the ovariectomy model does not parallel the menopausal transition in that during ovariectomy, circulating estrogens are depleted immediately. On the other hand, human menopause is preceded by peri-menopause, in which estrogen levels fluctuate appreciably. Imaging studies reveal that the changes in brain glucose utilization, neuropathology, cognitive alterations actually begin during peri-menopause in women^9-12^. In mice, in contrast to ovariectomy, a model of menopause that incorporates the peri-menopausal transition can be achieved by administering 4-vinylcyclohexene diepoxide (VCD). VCD accelerates ovarian failure while keeping ovaries intact^76,77^. Previous work using VCD has shown that VCD causes large estrogen fluctuations before ovarian failure, mimicking peri-menopause^77^. Using this model, our group and others have shown that accelerated ovarian failure leads to exacerbation of metabolic deficits^51,77^, neurovascular dysfunction^78^, and cognitive impairment^51^. We are currently testing the ability of DHED to rescue such deficits in this model of menopause.

In this study, we identified a new method of administering DHED long-term, via silastic capsule, with a dose of 1:10 DHED:cholesterol showing optimal results. We identified several new behavioral paradigms in which DHED exhibits beneficial effects in both young and middle-aged mice. Our data strongly supports the use of DHED to enhance cognitive performance in a rodent menopause model. By increasing estrogen exclusively in the brain, DHED represents a potential novel therapeutic strategy for cognitive decline in post-menopausal women, avoiding adverse peripheral effects of traditional estrogen replacement therapy.

## DECLARATIONS

### ETHICS APPROVAL AND CONSENT TO PARTICIPATE

Not applicable (no human subjects).

### CONSENT FOR PUBLICATION

Not applicable (no human subjects).

### AVAILABILITY OF DATA AND MATERIALS

The datasets acquired and/or analyzed during the current study are available from the corresponding author on reasonable request.

### COMPETING INTERESTS

The authors have no conflicts to disclose.

### FUNDING

This work was funded by an American Heart Association pre-doctoral award 908878 (AES), BrightFocus Foundation postdoctoral fellowship A2022001F (CAG), NINDS/NIA R01NS110749 (KLZ), NIA U01 AG072464 (KLZ), Alzheimer’s Association AARG-21-849204 (KLZ).

### AUTHORS’ CONTRIBUTIONS

KLZ obtained funding for the experiments. KLZ, AES, CAG, SBZS, KMH, and DZ designed the experiments. AES, CAG, HV, RMS, RDK, AS performed the animal work. AES and VN performed the RT-qPCR experiments. CAG, AS, VS, CAT, RS, KMB and OJG performed blinded analysis of behavior tests. AES and CAG analyzed the data. CAG and AES prepared the figures. KLZ, CAG, and AES interpreted the results. AES and CAG prepared the manuscript. KLZ edited the manuscript. All authors approved the final manuscript.

## ACKNOWLEDGEMENTS

None.

## Supplemental material

**Supplemental Figure 1.**
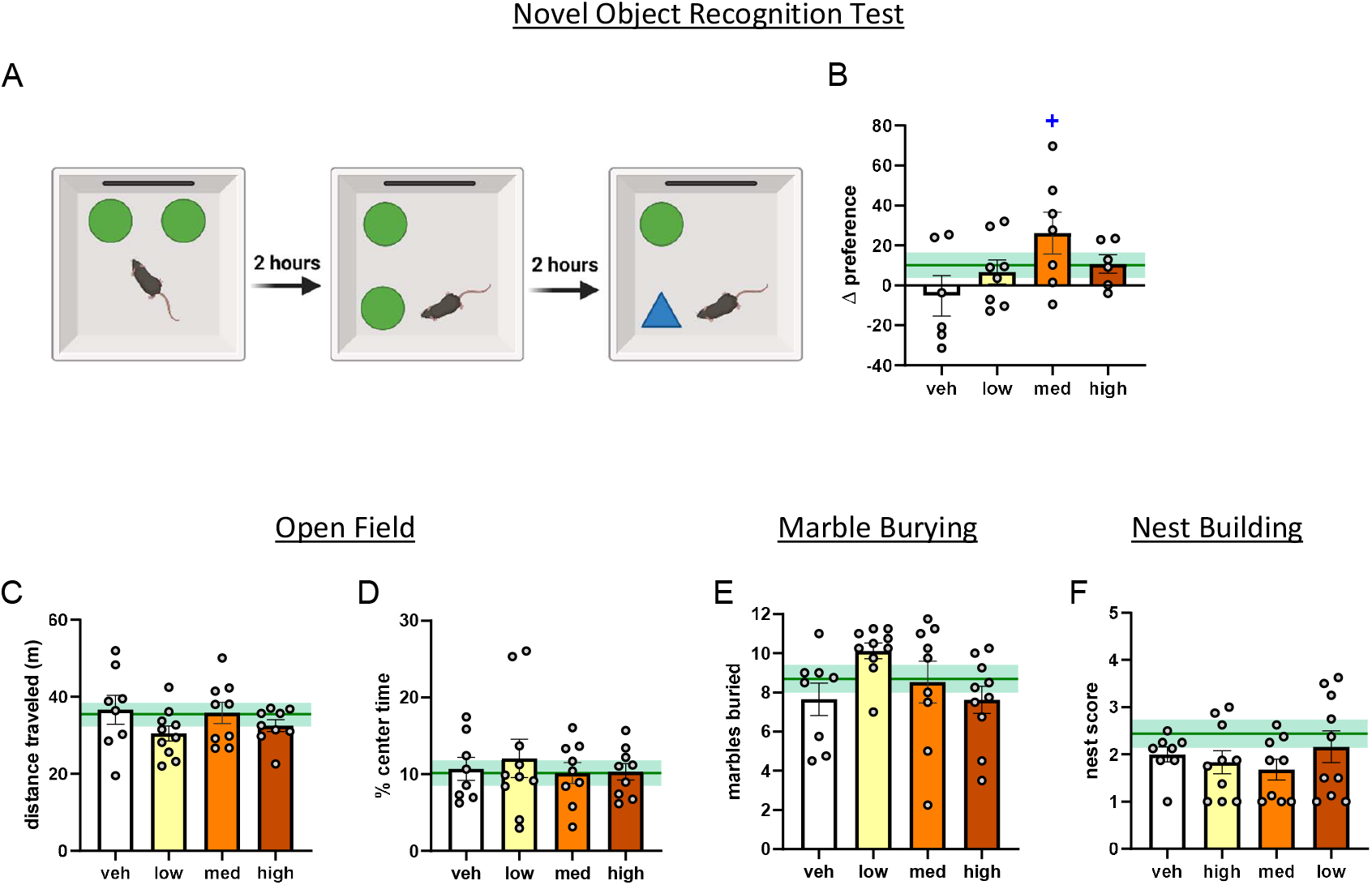
DHED does not alter anxiety-like behavior or activities of daily living. (A) Schematic of novel object recognition (created in BioRender) depicting the training trial, object place testing trial and novel object recognition testing trial. (B) Difference between % time spent with object in its new place during novel object recognition testing trial vs % time with replaced object during training trial. Open Field (C) distance traveled and (D) % time in the center of the open field arena. (E) Number of marbles buried during the marble burying test. (F) Nest scores from the nest building task. Data are presented as mean +SEM. +p<0.05, one-sample T-test vs chance (0). Data are represented as mean ±SEM. One-way ANOVA with Dunnett’s post hoc test vs vehicle *p<0.05. Gonadally intact, vehicle-treated age-matched reference group represented on bar graphs as mean (dark green line) ±SEM (light green shading). veh=vehicle; med=medium.

## Notes

### Competing Interest Statement

The authors have declared no competing interest.

## REFERENCES

1 Hill, K. The demography of menopause. Maturitas 23, 113–127, doi:10.1016/0378-5122(95)00968-x (1996).

2 Rocca, W. A., Grossardt, B. R. & Maraganore, D. M. The long-term effects of oophorectomy on cognitive and motor aging are age dependent. Neurodegener Dis 5, 257–260, doi:10.1159/000113718 (2008).

3 El Khoudary, S. R. et al. Menopause Transition and Cardiovascular Disease Risk: Implications for Timing of Early Prevention: A Scientific Statement From the American Heart Association. Circulation 142, e506–e532, doi:10.1161/cir.0000000000000912 (2020).

4 Stachowiak, G., Pertyński, T. & Pertyńska-Marczewska, M. Metabolic disorders in menopause. Prz Menopauzalny 14, 59–64, doi:10.5114/pm.2015.50000 (2015).

5 Lisabeth, L. & Bushnell, C. Stroke risk in women: the role of menopause and hormone therapy. Lancet Neurol 11, 82–91, doi:10.1016/s1474-4422(11)70269-1 (2012).

6 Cheng, G., Huang, C., Deng, H. & Wang, H. Diabetes as a risk factor for dementia and mild cognitive impairment: a meta-analysis of longitudinal studies. Intern Med J 42, 484–491, doi:10.1111/j.1445-5994.2012.02758.x (2012).

7 Pirimoglu, Z. M. et al. Glucose tolerance of premenopausal women after menopause due to surgical removal of ovaries. Climacteric 14, 453–457, doi:10.3109/13697137.2010.539723 (2011).

8 Slopien, R. et al. Menopause and diabetes: EMAS clinical guide. Maturitas 117, 6–10, doi:10.1016/j.maturitas.2018.08.009 (2018).

9 Mosconi, L. et al. Increased Alzheimer’s risk during the menopause transition: A 3-year longitudinal brain imaging study. PLoS One 13, e0207885, doi:10.1371/journal.pone.0207885 (2018).

10 Brinton, R. D., Yao, J., Yin, F., Mack, W. J. & Cadenas, E. Perimenopause as a neurological transition state. Nat Rev Endocrinol 11, 393–405, doi:10.1038/nrendo.2015.82 (2015).

11 Mosconi, L. et al. Sex differences in Alzheimer risk: Brain imaging of endocrine vs chronological aging. Neurology 89, 1382–1390, doi:10.1212/wnl.0000000000004425 (2017).

12 Rahman, A. et al. Sex-driven modifiers of Alzheimer risk: A multimodality brain imaging study. Neurology 95, e166–e178, doi:10.1212/wnl.0000000000009781 (2020).

13 Rocca, W. A. et al. Loss of Ovarian Hormones and Accelerated Somatic and Mental Aging. Physiology (Bethesda) 33, 374–383, doi:10.1152/physiol.00024.2018 (2018).

14 Georgakis, M. K. et al. Age at menopause and duration of reproductive period in association with dementia and cognitive function: A systematic review and meta-analysis. Psychoneuroendocrinology 73, 224–243, doi:10.1016/j.psyneuen.2016.08.003 (2016).

15 Coughlan, G. T. et al. Association of Age at Menopause and Hormone Therapy Use With Tau and beta-Amyloid Positron Emission Tomography. JAMA Neurol 80, 462–473, doi:10.1001/jamaneurol.2023.0455 (2023).

16 Gannon, O. J., Robison, L. S., Custozzo, A. J. & Zuloaga, K. L. Sex differences in risk factors for vascular contributions to cognitive impairment & dementia. Neurochemistry international 127, 38–55, doi:10.1016/j.neuint.2018.11.014 (2019).

17 Frick, K. M., Kim, J. & Koss, W. A. Estradiol and hippocampal memory in female and male rodents. Curr Opin Behav Sci 23, 65–74, doi:10.1016/j.cobeha.2018.03.011 (2018).

18 Jett, S. et al. Ovarian steroid hormones: A long overlooked but critical contributor to brain aging and Alzheimer’s disease. Front Aging Neurosci 14, 948219, doi:10.3389/fnagi.2022.948219 (2022).

19 Cheng, C. M., Cohen, M., Wang, J. & Bondy, C. A. Estrogen augments glucose transporter and IGF1 expression in primate cerebral cortex. Faseb j 15, 907–915, doi:10.1096/fj.00-0398com (2001).

20 Shi, J. & Simpkins, J. W. 17 beta-Estradiol modulation of glucose transporter 1 expression in blood-brain barrier. Am J Physiol 272, E1016–1022, doi:10.1152/ajpendo.1997.272.6.E1016 (1997).

21 Maggioli, E. et al. Estrogen protects the blood-brain barrier from inflammation-induced disruption and increased lymphocyte trafficking. Brain Behav Immun 51, 212–222, doi:10.1016/j.bbi.2015.08.020 (2016).

22 Villa, A., Rizzi, N., Vegeto, E., Ciana, P. & Maggi, A. Estrogen accelerates the resolution of inflammation in macrophagic cells. Sci Rep 5, 15224, doi:10.1038/srep15224 (2015).

23 Lu, Y. et al. Neuron-Derived Estrogen Regulates Synaptic Plasticity and Memory. J Neurosci 39, 2792–2809, doi:10.1523/jneurosci.1970-18.2019 (2019).

24 Frick, K. M. Molecular mechanisms underlying the memory-enhancing effects of estradiol. Horm Behav 74, 4–18, doi:10.1016/j.yhbeh.2015.05.001 (2015).

25 Sheppard, P. A. S., Choleris, E. & Galea, L. A. M. Structural plasticity of the hippocampus in response to estrogens in female rodents. Mol Brain 12, 22, doi:10.1186/s13041-019-0442-7 (2019).

26 Galea, L. A. M., Frick, K. M., Hampson, E., Sohrabji, F. & Choleris, E. Why estrogens matter for behavior and brain health. Neurosci Biobehav Rev 76, 363–379, doi:10.1016/j.neubiorev.2016.03.024 (2017).

27 Sierra, A., Gottfried-Blackmore, A., Milner, T. A., McEwen, B. S. & Bulloch, K. Steroid hormone receptor expression and function in microglia. Glia 56, 659–674, doi:10.1002/glia.20644 (2008).

28 Arevalo, M. A., Santos-Galindo, M., Bellini, M. J., Azcoitia, I. & Garcia-Segura, L. M. Actions of estrogens on glial cells: Implications for neuroprotection. Biochim Biophys Acta 1800, 1106–1112, doi:10.1016/j.bbagen.2009.10.002 (2010).

29 Azcoitia, I., Sierra, A. & Garcia-Segura, L. M. Localization of estrogen receptor beta-immunoreactivity in astrocytes of the adult rat brain. Glia 26, 260–267 (1999).

30 Jung-Testas, I. & Baulieu, E. E. Steroid hormone receptors and steroid action in rat glial cells of the central and peripheral nervous system. J Steroid Biochem Mol Biol 65, 243–251, doi:10.1016/s0960-0760(97)00191-x (1998).

31 Zuloaga, D. G., Zuloaga, K. L., Hinds, L. R., Carbone, D. L. & Handa, R. J. Estrogen receptor beta expression in the mouse forebrain: age and sex differences. J Comp Neurol 522, 358–371, doi:10.1002/cne.23400 (2014).

32 Zuloaga, K. L., Swift, S. N., Gonzales, R. J., Wu, T. J. & Handa, R. J. The androgen metabolite, 5alpha-androstane-3beta,17beta-diol, decreases cytokine-induced cyclooxygenase-2, vascular cell adhesion molecule-1 expression, and P-glycoprotein expression in male human brain microvascular endothelial cells. Endocrinology 153, 5949–5960, doi:10.1210/en.2012-1316 (2012).

33 Milner, T. A. et al. Ultrastructural localization of estrogen receptor beta immunoreactivity in the rat hippocampal formation. J Comp Neurol 491, 81–95, doi:10.1002/cne.20724 (2005).

34 Wilson, M. E., Westberry, J. M. & Trout, A. L. Estrogen receptor-alpha gene expression in the cortex: sex differences during development and in adulthood. Horm Behav 59, 353–357, doi:10.1016/j.yhbeh.2010.08.004 (2011).

35 Sárvári, M. et al. Hippocampal Gene Expression Is Highly Responsive to Estradiol Replacement in Middle-Aged Female Rats. Endocrinology 156, 2632–2645, doi:10.1210/en.2015-1109 (2015).

36 Dean, B. & Gogos, A. The impact of ovariectomy and chronic estrogen treatment on gene expression in the rat cortex: Implications for psychiatric disorders. Psychoneuroendocrinology 127, 105192, doi:10.1016/j.psyneuen.2021.105192 (2021).

37 Fox, M., Berzuini, C. & Knapp, L. A. Cumulative estrogen exposure, number of menstrual cycles, and Alzheimer’s risk in a cohort of British women. Psychoneuroendocrinology 38, 2973–2982, doi:10.1016/j.psyneuen.2013.08.005 (2013).

38 Matyi, J. M., Rattinger, G. B., Schwartz, S., Buhusi, M. & Tschanz, J. T. Lifetime estrogen exposure and cognition in late life: the Cache County Study. Menopause 26, 1366–1374, doi:10.1097/gme.0000000000001405 (2019).

39 Rossouw, J. E. et al. Risks and benefits of estrogen plus progestin in healthy postmenopausal women: principal results From the Women’s Health Initiative randomized controlled trial. Jama 288, 321–333 (2002).

40 Henderson, V. W. et al. Estrogen for Alzheimer’s disease in women: randomized, double-blind, placebo-controlled trial. Neurology 54, 295–301, doi:10.1212/wnl.54.2.295 (2000).

41 Wang, P. N. et al. Effects of estrogen on cognition, mood, and cerebral blood flow in AD: a controlled study. Neurology 54, 2061–2066, doi:10.1212/wnl.54.11.2061 (2000).

42 Miller, V. M. & Harman, S. M. An update on hormone therapy in postmenopausal women: mini-review for the basic scientist. American journal of physiology. Heart and circulatory physiology 313, H1013–h1021, doi:10.1152/ajpheart.00383.2017 (2017).

43 Zandi, P. P. et al. Hormone replacement therapy and incidence of Alzheimer disease in older women: the Cache County Study. Jama 288, 2123–2129 (2002).

44 Marjoribanks, J., Farquhar, C., Roberts, H., Lethaby, A. & Lee, J. Long-term hormone therapy for perimenopausal and postmenopausal women. Cochrane Database Syst Rev 1, Cd004143, doi:10.1002/14651858.CD004143.pub5 (2017).

45 Prokai, L. et al. The prodrug DHED selectively delivers 17β-estradiol to the brain for treating estrogen-responsive disorders. Sci Transl Med 7, 297ra113, doi:10.1126/scitranslmed.aab1290 (2015).

46 Merchenthaler, I. et al. Treatment with an orally bioavailable prodrug of 17β-estradiol alleviates hot flushes without hormonal effects in the periphery. Sci Rep 6, 30721, doi:10.1038/srep30721 (2016).

47 Prokai-Tatrai, K., Nguyen, V. & Prokai, L. 10β,17α-Dihydroxyestra-1,4-dien-3-one: A Bioprecursor Prodrug Preferentially Producing 17α-Estradiol in the Brain for Targeted Neurotherapy. ACS Chem Neurosci 9, 2528–2533, doi:10.1021/acschemneuro.8b00184 (2018).

48 Ronan, L. et al. Obesity associated with increased brain age from midlife. Neurobiol Aging 47, 63–70, doi:10.1016/j.neurobiolaging.2016.07.010 (2016).

49 Salinero, A. E. et al. Sex-specific effects of high-fat diet on cognitive impairment in a mouse model of VCID. FASEB journal : official publication of the Federation of American Societies for Experimental Biology 34, 15108–15122, doi:10.1096/fj.202000085R (2020).

50 Gannon, O. J. et al. High-fat diet exacerbates cognitive decline in mouse models of Alzheimer’s disease and mixed dementia in a sex-dependent manner. Journal of neuroinflammation 19, 110, doi:10.1186/s12974-022-02466-2 (2022).

51 Gannon, O. J. et al. Menopause causes metabolic and cognitive impairments in a chronic cerebral hypoperfusion model of vascular contributions to cognitive impairment and dementia. Biology of sex differences 14, 34, doi:10.1186/s13293-023-00518-7 (2023).

52 Deacon, R. M. Assessing nest building in mice. Nat Protoc 1, 1117–1119, doi:10.1038/nprot.2006.170 (2006).

53 Salinero, A. E., Anderson, B. M. & Zuloaga, K. L. Sex differences in the metabolic effects of diet-induced obesity vary by age of onset. Int J Obes (Lond) 42, 1088–1091, doi:10.1038/s41366-018-0023-3 (2018).

54 Yang, H. R., Tu, T. H., Jeong, D. Y., Yang, S. & Kim, J. G. Obesity induced by estrogen deficiency is associated with hypothalamic inflammation. Biochem Biophys Rep 23, 100794, doi:10.1016/j.bbrep.2020.100794 (2020).

55 Stefanska, A., Bergmann, K. & Sypniewska, G. Metabolic Syndrome and Menopause: Pathophysiology, Clinical and Diagnostic Significance. Adv Clin Chem 72, 1–75, doi:10.1016/bs.acc.2015.07.001 (2015).

56 Pu, D., Tan, R., Yu, Q. & Wu, J. Metabolic syndrome in menopause and associated factors: a meta-analysis. Climacteric 20, 583–591, doi:10.1080/13697137.2017.1386649 (2017).

57 Manrique-Acevedo, C., Chinnakotla, B., Padilla, J., Martinez-Lemus, L. A. & Gozal, D. Obesity and cardiovascular disease in women. Int J Obes (Lond) 44, 1210–1226, doi:10.1038/s41366-020-0548-0 (2020).

58 Foster, T. C. Role of estrogen receptor alpha and beta expression and signaling on cognitive function during aging. Hippocampus 22, 656–669, doi:10.1002/hipo.20935 (2012).

59 Bean, L. A., Ianov, L. & Foster, T. C. Estrogen receptors, the hippocampus, and memory. Neuroscientist 20, 534–545, doi:10.1177/1073858413519865 (2014).

60 Tschiffely, A. E., Schuh, R. A., Prokai-Tatrai, K., Prokai, L. & Ottinger, M. A. A comparative evaluation of treatments with 17β-estradiol and its brain-selective prodrug in a double-transgenic mouse model of Alzheimer’s disease. Horm Behav 83, 39–44, doi:10.1016/j.yhbeh.2016.05.009 (2016).

61 Markowska, A. L. & Savonenko, A. V. Effectiveness of estrogen replacement in restoration of cognitive function after long-term estrogen withdrawal in aging rats. J Neurosci 22, 10985–10995, doi:10.1523/jneurosci.22-24-10985.2002 (2002).

62 Acharya, K. D. et al. Estradiol-mediated protection against high-fat diet induced anxiety and obesity is associated with changes in the gut microbiota in female mice. Sci Rep 13, 4776, doi:10.1038/s41598-023-31783-6 (2023).

63 Vigil, P., Meléndez, J., Petkovic, G. & Del Río, J. P. The importance of estradiol for body weight regulation in women. Front Endocrinol (Lausanne) 13, 951186, doi:10.3389/fendo.2022.951186 (2022).

64 Hrupka, B. J., Smith, G. P. & Geary, N. Hypothalamic implants of dilute estradiol fail to reduce feeding in ovariectomized rats. Physiol Behav 77, 233–241, doi:10.1016/s0031-9384(02)00857-0 (2002).

65 Butera, P. C. & Beikirch, R. J. Central implants of diluted estradiol: independent effects on ingestive and reproductive behaviors of ovariectomized rats. Brain Res 491, 266–273, doi:10.1016/0006-8993(89)90062-0 (1989).

66 Frick, K. M. & Berger-Sweeney, J. Spatial reference memory and neocortical neurochemistry vary with the estrous cycle in C57BL/6 mice. Behav Neurosci 115, 229–237, doi:10.1037/0735-7044.115.1.229 (2001).

67 Frye, C. A. Estrus-associated decrements in a water maze task are limited to acquisition. Physiol Behav 57, 5–14, doi:10.1016/0031-9384(94)00197-d (1995).

68 Tuscher, J. J., Fortress, A. M., Kim, J. & Frick, K. M. Regulation of object recognition and object placement by ovarian sex steroid hormones. Behav Brain Res 285, 140–157, doi:10.1016/j.bbr.2014.08.001 (2015).

69 Luine, V. N., Jacome, L. F. & Maclusky, N. J. Rapid enhancement of visual and place memory by estrogens in rats. Endocrinology 144, 2836–2844, doi:10.1210/en.2003-0004 (2003).

70 Frye, C. A., Duffy, C. K. & Walf, A. A. Estrogens and progestins enhance spatial learning of intact and ovariectomized rats in the object placement task. Neurobiol Learn Mem 88, 208–216, doi:10.1016/j.nlm.2007.04.003 (2007).

71 Tuscher, J. J. et al. Inhibition of local estrogen synthesis in the hippocampus impairs hippocampal memory consolidation in ovariectomized female mice. Horm Behav 83, 60–67, doi:10.1016/j.yhbeh.2016.05.001 (2016).

72 Kim, J. & Frick, K. M. Distinct effects of estrogen receptor antagonism on object recognition and spatial memory consolidation in ovariectomized mice. Psychoneuroendocrinology 85, 110–114, doi:10.1016/j.psyneuen.2017.08.013 (2017).

73 Zhao, L. & Brinton, R. D. Estrogen receptor alpha and beta differentially regulate intracellular Ca(2+) dynamics leading to ERK phosphorylation and estrogen neuroprotection in hippocampal neurons. Brain Res 1172, 48–59, doi:10.1016/j.brainres.2007.06.092 (2007).

74 Vedder, L. C., Bredemann, T. M. & McMahon, L. L. Estradiol replacement extends the window of opportunity for hippocampal function. Neurobiol Aging 35, 2183–2192, doi:10.1016/j.neurobiolaging.2014.04.004 (2014).

75 Suzuki, S. et al. Timing of estrogen therapy after ovariectomy dictates the efficacy of its neuroprotective and antiinflammatory actions. Proc Natl Acad Sci U S A 104, 6013–6018, doi:10.1073/pnas.0610394104 (2007).

76 Kappeler, C. J. & Hoyer, P. B. 4-vinylcyclohexene diepoxide: a model chemical for ovotoxicity. Syst Biol Reprod Med 58, 57–62, doi:10.3109/19396368.2011.648820 (2012).

77 Brooks, H. L., Pollow, D. P. & Hoyer, P. B. The VCD Mouse Model of Menopause and Perimenopause for the Study of Sex Differences in Cardiovascular Disease and the Metabolic Syndrome. Physiology (Bethesda) 31, 250–257, doi:10.1152/physiol.00057.2014 (2016).

78 Blackwell, J. A. et al. Cerebral arteriolar and neurovascular dysfunction after chemically induced menopause in mice. Am J Physiol Heart Circ Physiol 323, H845–h860, doi:10.1152/ajpheart.00276.2022 (2022).

